# Genomic approaches to build de novo elite breeding gene pools from locally-adapted landraces

**DOI:** 10.1101/2025.06.30.662425

**Authors:** Safiétou Tooli Fall, Alexander Kena, Brian R. Rice, Ghislain Kanfany, Cyril Diatta, Ndjido A. Kane, Allan K. Fritz, Geoffrey P. Morris

## Abstract

Many nascent breeding programs aim to achieve genetic gain by crossing locally elite germplasm, but a lack of systematic approaches to develop elite gene pools from locally-adapted varieties hinders their progress. Motivated by the observation of undesirable transgressive segregation in presumed elite crosses in Senegalese cereal breeding programs, we designed approaches for de novo development of elite gene pools from locally-adapted landrace-derived germplasm. We first define two types of “elite” germplasm: *cis*-elite, phenotypically similar and genetically homogeneous for locally-adapted traits (“acquired traits”); versus *trans*-elite, phenotypically similar, but genetically heterogeneous for acquired traits. Next, we defined two genomic approaches for de novo inference of elite gene pools: population-based genotypic inference (PGI) and QTL-based genotypic inference (QGI), and compared to a family-based phenotypic inference (FPI) approach. Using simulations that trace the evolution from locally-adapted landraces to elite breeding lines, we evaluate the effectiveness of these strategies in nascent forward breeding programs. QGI accurately and cost-effectively identifies both *cis-* and *trans*-elite pairs, regardless of the underlying trait architecture, while PGI is less sensitive when trait architecture is oligogenic. Over ten cycles of phenotypic recurrent selection, programs based on *cis*-elite crosses consistently outperformed those based on *trans*-elite crosses for genetic gain. The findings highlight the value of trait genetic architecture knowledge for elite gene pool development and provide a practical roadmap for elite germplasm development in modernizing breeding programs.

## INTRODUCTION

Intercrossing elite lines to develop improved cultivars is at the core of modern plant breeding (Rutkoski, 2019). Despite widespread use of “elite” in plant breeding literature, however, there remains no universally accepted definition of *elite*, nor systematic guidance on how to develop *elite* germplasm (Acquaah, 2012; Bernardo, 2020; Falconer, 1996; Falk, 2010). The term is variously applied to denote superior phenotypic performance, the potential to produce superior progeny, and/or genetic similarity within breeding populations (Table S1). Advanced breeding programs, such as public-sector barley programs or private-sector corn breeding programs in North America and Europe, have typically managed narrow elite gene pools (Bornowski et al., 2021; Rasmusson & Phillips, 1997). By contrast, emerging breeding programs often begin with much more diverse local andraces that are well-appreciated by local farmers and consumers, but the lack of clearly defined elite gene pools makes it difficult to design elite-by-elite crosses (Kane et al., 2022). Conversely, attempts to import “elite” germplasm from established breeding programs in different geographies has often failed because that germplasm is maladapted to local environments and cultural needs of smallholder farmers (Walker & Alwang, 2015). Therefore, new approaches are needed to develop elite germplasm de novo from locally-adapted landrace germplasm.

Genetic gain requires deliberate population improvement strategies that balance stabilizing selection to maintain favorable traits and directional selection to drive progress toward breeding goals (Falconer, 1996; Hill, 2010). Stabilizing selection helps preserve adaptive genetic variation by maintaining key allele frequencies, while directional selection promotes the accumulation of favorable alleles to improve target traits (Walsh & Lynch, 2018a). The interplay between these selective forces underlies the genetic architecture of elite germplasm and influences long term response to selection (Hill, 2010; Walsh & Lynch, 2018a). Breeding programs that manage this balance effectively sustain additive genetic variance, which according to the breeder’s equation, is critical for maximizing genetic gain (Lush, 1943).

Landrace origins are shaped by a combination of farmer selection preferences, local environmental pressures, and seed exchange networks, which can promote genetic heterogeneity through diverse adaptation or lead to homogeneity under isolation or intensive selection (Nelimor et al., 2020; Rattunde et al., 2021). In crops such as sorghum, millet, and cowpea, this diversity is a critical resource, but it complicates the definition and development of elite germplasm. Phenotypic similarity among lines may mask underlying genetic heterogeneity (Brown, 1989; Crossa et al., 2017), and individual-based selection risks overlooking this cryptic variation (R & B, 1981). In contrast, established programs have often used family-based selection to capture hidden structure and stabilize elite populations (Bernardo, 2020; Comstock & Robinson, 1948). These contrasting strategies highlight the need for emerging programs to adopt more deliberate frameworks that integrate diversity and structure in the development of elite gene pools.

Population genetic theory on parallel evolution is one promising area to inform frameworks for elite gene pool development. When distinct lineages from related founder populations are exposed to similar environmental pressures, they often evolve toward similar phenotypes, a common outcome in evolution known as parallel evolution (Wood et al., 2005). The evolutionary processes leading to genomic parallelism when multiple lineages independently evolve toward a similar phenotype in similar environments (Bohutínská et al., 2021). Individuals from different closely related founding populations adapt in similar ways under the same conditions with different genetics (Blount et al., 2018; Whiting et al., 2022). In addition, the pressure and direction of selection and genetic drift applied to individual quantitative trait loci (QTL) can give rise to alleles with an antagonistic effect for the trait, and thus increase the extent and frequency of transgressive segregation from crosses in domesticated populations (Dickinson et al., 2003). While parallel evolution has been studied in domestication and local adaptation, its implications for crop breeding remain largely underexplored.

Here, we present a framework for de novo elite gene pool development in emerging breeding programs through the use of molecular markers. Our objective was to identify accurate, efficient, and cost-effective strategies for inferring elite genetic relationships within diverse germplasm pools to optimize crossing decisions and accelerate genetic gain. To this end, we compared a phenotypic inference approach, family-based phenotypic inference (FPI), with two new genomic inference approaches, population-based genotypic inference (PGI) and QTL-based genotypic inference (QGI), evaluating their effectiveness across traits with differing genetic architectures. Our findings demonstrate that genotypic inference approaches can be successfully applied to establish elite gene pools, with particular strengths depending on the underlying genetic basis of target traits. These results highlight an opportunity to deploy genetic knowledge for de novo development of elite gene pools, offering practical guidance for emerging breeding programs aiming to balance eliteness with genetic diversity.

## RESULTS

### Unexpected transgressive segregation from presumed elite breeding crosses

To assess the status of eliteness within the germplasm of emerging breeding programs, we analyzed progenies from recent elite-by-elite crosses made by breeders, along with effectiveness of ongoing trait introgression efforts. The motivating example for this study was the observation of transgressive segregation in putative elite-by-elite Senegalese sorghum and pearl millet breeding programs (Figures 1A–D). In pearl millet, a cross between the high-yielding, early-flowering variety Souna 3 and Thialack 2, valued for its long panicles, resulted in only 33% of F_5_ progenies flowering within the acceptable range (Figures 1A), undermining expectations for combining favorable traits (Figures 1C). A similar outcome occurred with a sorghum cross between two late-flowering lines, Nguinthe and Grinkan, intended to enhance panicle size (Figures 1B and 1D). In both cases, the broad segregation for flowering time reduced the number of candidates for advancement, and therefore the effective population size (Figures 1C–D).

**Figure 1.**
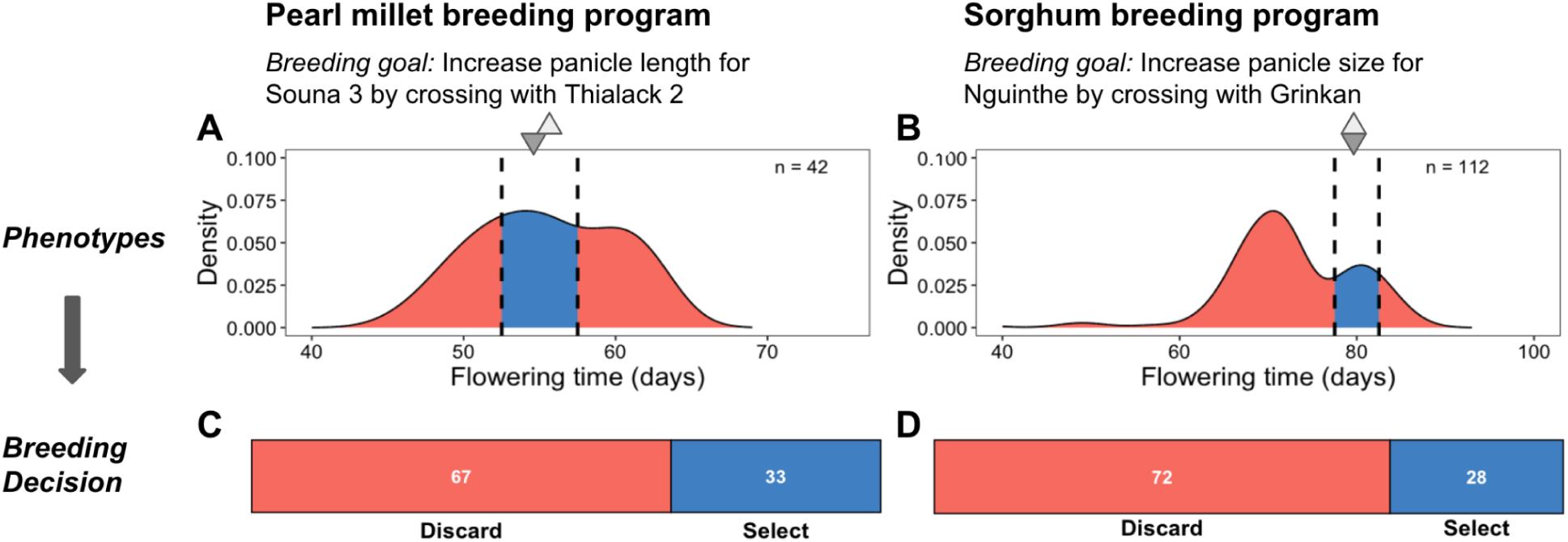
Undesirable transgressive segregation for an acquired trait in progeny from presumed elite-by-elite crosses substantially reduces the number of candidates for selection. Phenotypic distribution of flowering time in progeny from presumed elite-by-elite crosses in millet (A; F_5_ progeny) and sorghum (B; F_7_ progeny) breeding programs in Senegal. For pearl millet, Souna 3 was crossed with Thialack 2 to improve panicle length while maintaining earliness and high yield. For sorghum, Nguinthe was crossed with Grinkan to improve panicle size while maintaining lateness. Flowering time is an acquired trait in these programs, since the parents (triangles) are within the acceptable range of phenotypes (black dotted lines) according to the breeding product profiles. For both crops, substantial transgressive segregation was observed, with few progeny within the acceptable range (blue shading in A and B), and most progeny falling outside (red shading in A and B). Accordingly, the percentage progeny of discarded (red in C and D) versus selected (blue in C and D) progeny was high, with only 33% and 28% of the progeny for pearl millet and sorghum, respectively, selected as acceptable with respect to flowering time.

### Conceptual model for genetic heterogeneity in elite germplasm: *cis-* versus *trans*-eliteness

In many emerging breeding programs particularly across sub-Saharan Africa early efforts over the past century focused on collecting and stabilizing locally adapted landraces into pure line varieties (Figure 2A). These presumed elite lines were developed to consolidate key adaptive traits, such as flowering time, while enhancing agronomic performance traits like yield and disease resistance (Figure 2A). Traits for which an acceptable phenotypic value has already been captured during this process are referred to here as “acquired traits” (under stabilizing selection). By contrast, “desired traits” (under directional selection) are those for which breeding efforts aim to change the phenotypic value, often to meet farmer preferences or evolving environmental demands. Crosses are often made based on phenotypic similarity without accounting for underlying genetic relatedness or genotypic complementarity. We hypothesize that excessive transgressive segregation observed in crosses of similar parent lines may be explained by cryptic genetic heterogeneity shaped by parallel evolution in isolated breeding efforts (Figure 2B–C).

**Figure 2.**
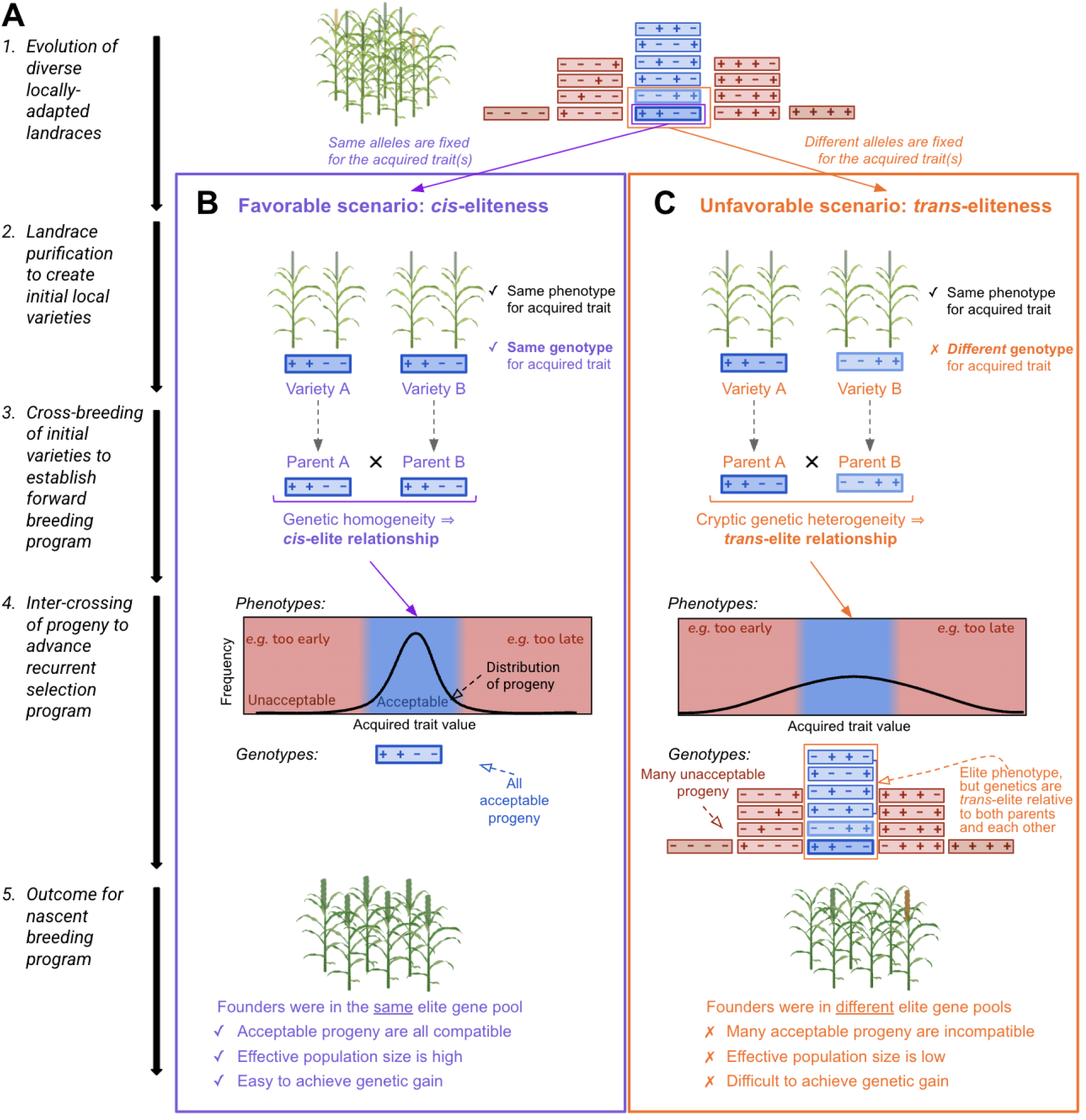
Evolutionary scenarios that lead to *cis-* versus *trans*-elite relationships in nascent breeding programs. (A) Schematic overview of the breeding trajectory, starting from landrace collection and parallel adaptation (Step 1), followed by landrace purification (Step 2), initial crossing of locally-elite lines (Step 3), intercrossing in recurrent breeding programs (Step 4), and culminating in the outcomes observed in nascent breeding programs (Step 5). Two distinct evolutionary scenarios are considered at each step: (B) *cis*-eliteness, characterized by genetic homogeneity among elite parents, and (C) *trans*-eliteness, characterized by cryptic genetic heterogeneity. Each rectangle represents the diploid genotype of a line, with “+” and “–” indicating favorable and unfavorable alleles, respectively, across four representative QTL for an acquired traits. (Heterozygosity is ignored for simplicity). In the *cis*-elite scenario (B), genetically homogeneous elite parents produce progeny with a narrow phenotypic distribution, where most offspring fall within an acceptable range. By contrast, the *trans*-elite scenario involves genetically heterogeneous elite parents, resulting in a broader phenotypic distribution, with many progeny falling outside the acceptable range and being discarded.

Thus, we propose that two contrasting types of elite relationships may exist: *cis*-versus *trans*-elite. In the *cis*-elite cross scenario, repeated selection and genetic drift within locally adapted germplasm may result in elite lines that share similar genotypes at acquired traits (Figure 2B). Crosses between such genetically homogeneous lines typically produce narrow phenotypic distributions, with most progeny mostly resembling the parents in both trait phenotype and genetic makeup. This facilitates efficient selection and helps maintain a robust effective population size (Figure 2B). By contrast, the *trans*-elite cross scenario would occur after a history of parallel adaptation of populations with the same phenotype, but different genotypes at acquired traits. Here, elite lines derive from different ancestral lineages and carry distinct haplotypes, with alleles in repulsion phase (Figure 2C). Crosses between such divergent lines can lead to extensive transgressive segregation due to the recombination of incompatible or novel allelic combinations (Figure 2C). While this genetic novelty may occasionally yield superior individuals for desired traits, it more often results in progeny with unacceptable values for acquired traits. This would lower the proportion of acceptable progeny, reducing effective population size, and therefore, genetic gain.

### Proposed approaches to infer *cis*-versus *trans*-elite relationships

We propose three approaches to infer elite line relationships, each grounded in a distinct genetic rationale and suited to varying resource capacities:

- *Family-based Phenotypic Inference*: FPI would involve crossing elite lines in a diallel or partial diallel design and phenotypically evaluating the resulting segregating progeny (e.g., F_2_ to F_5_ generations) (Figure 3A). The theoretical expectation is that *cis*-elite pairs, which share similar haplotypes, will produce narrow phenotypic distributions, while *trans*-elite pairs will yield broader segregation due to greater allelic divergence. Since this approach requires only classical breeding techniques (crossing, phenotyping, and selection) it may have been how (intentionally or unintentionally) classical breeders in the twentieth century developed elite gene pools. This approach provides inferences without genomic or molecular tools but is time-consuming and resource-intensive.
- *Population-based Genotypic Inference:* PGI would leverage genome-wide marker data to infer population structure among elite lines (Figure 3B). We applied Discriminant Analysis of Principal Components (DAPC), a multivariate method that clusters individuals based on genetic variation without assuming Hardy-Weinberg equilibrium. DAPC maximizes between group variation while minimizing within group variance, allowing the identification of genetically homogeneous clusters. The rationale is that elite lines belonging to the same genetic cluster (i.e., same subpopulation) are more likely to share ancestral haplotypes, especially in genomic regions under selection. Such lines are thus inferred as having a *cis*-elite relationship, reducing the likelihood of unexpected segregation when crossed.
- *QTL-based Genotypic Inference:* QGI would use genetic similarity within known QTL that underlie acquired traits that are essential for acceptability (e.g. flowering time, plant height, or grain color in cereals) (Figure 3C). Identity-by-state (IBS) matrices are used to compare elite lines based on shared allelic states at locus-specific SNPs within trait-associated QTLs. This approach provides a more precise assessment of genetic compatibility by directly examining variation at underlying loci rather than genome-wide background. Elite lines that share high IBS values at relevant QTLs are considered *cis*-elite pairs with respect to the studied trait, whereas those with divergent alleles enabled allelic recombination increase the risk for transgressive segregation and therefore inferred *trans*-elite pairs. QGI is theoretically the most targeted and informative approach, but it depends on prior knowledge of QTL or quantitative trait nucleotides (QTNs) and access to high-quality genotypic data.

**Figure 3.**
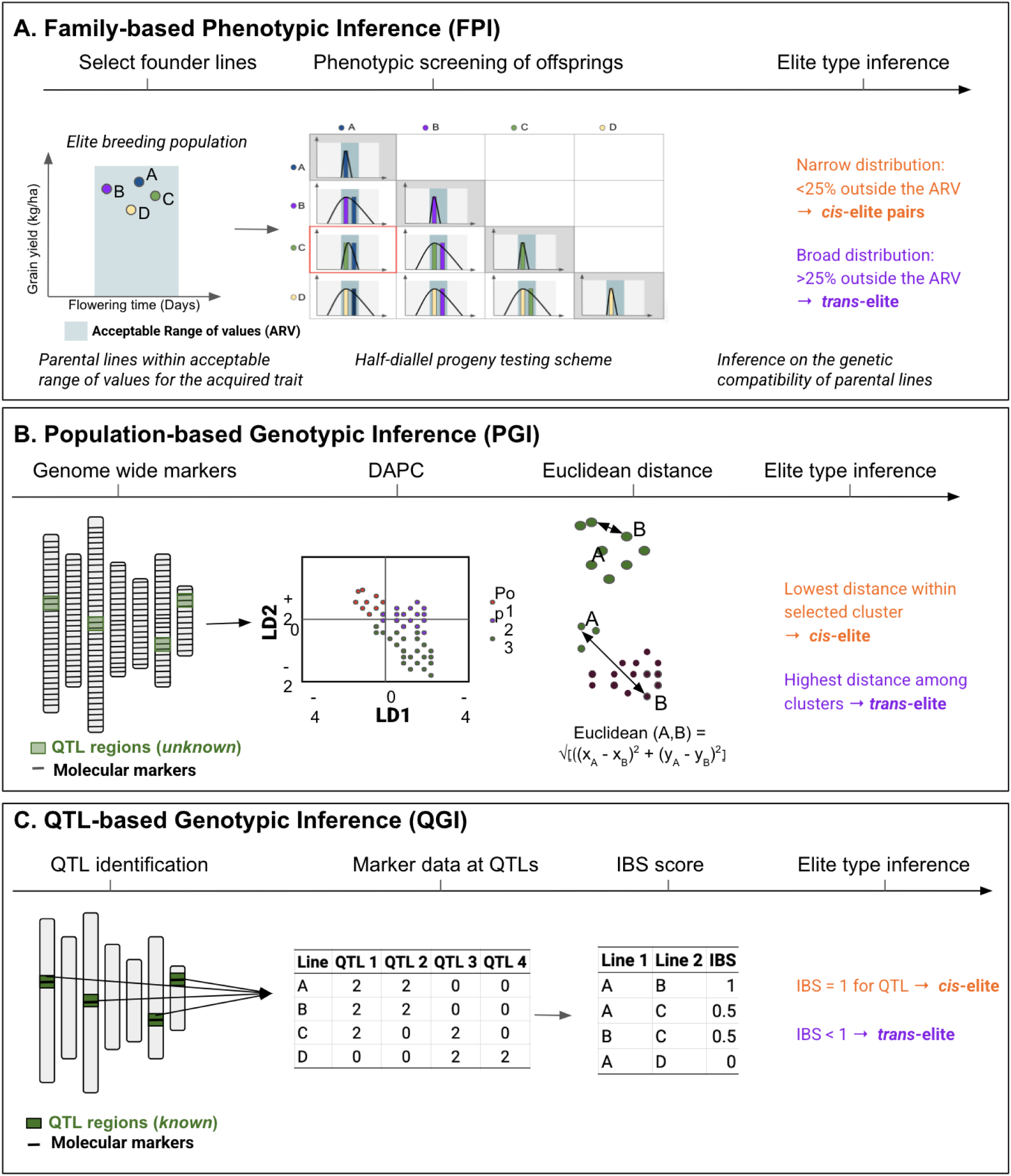
Schematic overview of approaches to inference *cis-* and *trans-* eliteness. (A) *Family-based phenotypic inference:* Elite lines were crossed using a half-diallel mating design. The *cis*- or *trans*-elite relationships were inferred from the phenotypic distribution of F_2_ progeny, and the accuracy of these inferences was evaluated using F_5_ progeny. Narrow phenotypic distributions were indicative of *cis*-eliteness, while broader distributions suggested *trans*-eliteness. (B) *Population-based genotypic inference (PGI).* Whole-genome marker data were used to perform Discriminant Analysis of Principal Components (DAPC) and to calculate pairwise Euclidean genetic distances. Lineages from the same subpopulation, showing low genetic distances, were inferred to have *cis*-elite relationships, while those from different subpopulations, showing higher distances, were inferred to have *trans*-elite relationships. (C) *QTL-based genotypic inference (QGI).* Identity-by-state (IBS) scores were calculated using markers located within known QTL. A score of IBS = 1 was interpreted as evidence of *cis*-eliteness, while IBS < 1 was interpreted as *trans*-eliteness.

To evaluate the potential effectiveness of these approaches we simulated the steps in the evolution of nascent breeding programs (Figure 2) and applied each of the approaches (Figure 3) to the simulated breeding programs (Figure 4; see Materials and Methods for details). These steps included: (1) creating diverse, locally-adapted landraces and defining the trait (Figure 4A), (2) purifying landraces to develop initial local varieties (Figure 4B), (3) cross-breeding initial varieties to establish a forward breeding program (Figure 4C), (4) inter-crossing progeny of *cis*-vs. *trans*-elite relationships to initiate a recurrent selection program (Figure 4D), and (5) evaluating outcomes for the nascent breeding program (Figure 4E).

**Figure 4.**
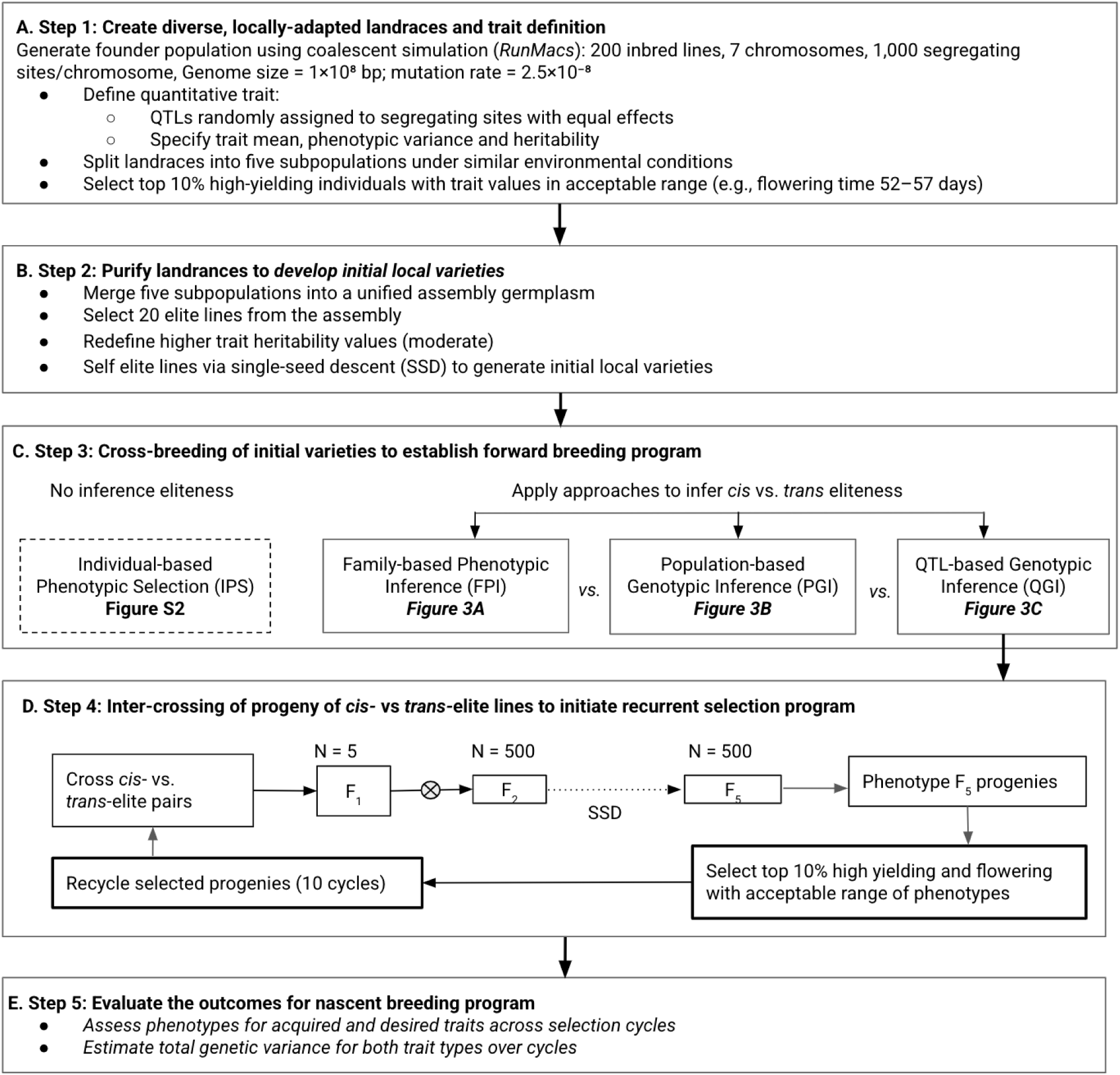
Simulation workflow for de novo creation, deployment, and evaluation of elite gene pools in nascent breeding programs. We conducted a five-step simulation process: (A) Generation of diverse, locally adapted landraces with defined trait architecture; (B) Purification and development of initial local varieties; (C) Cross-breeding to initiate a forward breeding program; (D) Intercrossing progeny with *cis*- or *trans*-elite relationships to launch phenotypic recurrent selection; (E) Evaluate the outcomes for nascent breeding program.

### Family-based phenotypic inference is simple but limited to simple acquired trait architecture

We evaluated the effectiveness of FPI in distinguishing *cis*- and *trans*-elite relationships by analyzing F_2_ progeny distributions and validating predictions against F_5_ outcomes under simulated genetic architectures involving 2, 4, or 20 QTLs (Figures 5A–C). However, across all QTL scenarios, FPI showed no consistent phenotypic pattern reliably distinguishing the two elite types (Figure 5A for QTL = 2, and Figure 5B for QTL = 4, and Figure 5C for QTL = 20). To apply the approach, we introduced a breeder-defined threshold: crosses were classified as *trans*-elite if more than 75% of F_2_ individuals fell outside the predefined acceptable range of phenotypes. Under this criterion, FPI correctly identified elite type in 70% of cases when 4 QTLs were involved, suggesting that the approach can be moderately effective when clear trait thresholds are established. Despite its simplicity and intuitive appeal, performance of FPI declined under more complex genetic architectures, highlighting its limitations as trait architecture complexity increases. Thus, while FPI offers a straightforward framework for elite-type inference, its accuracy is contingent on both trait complexity and specific phenotypic thresholds.

**Figure 5.**
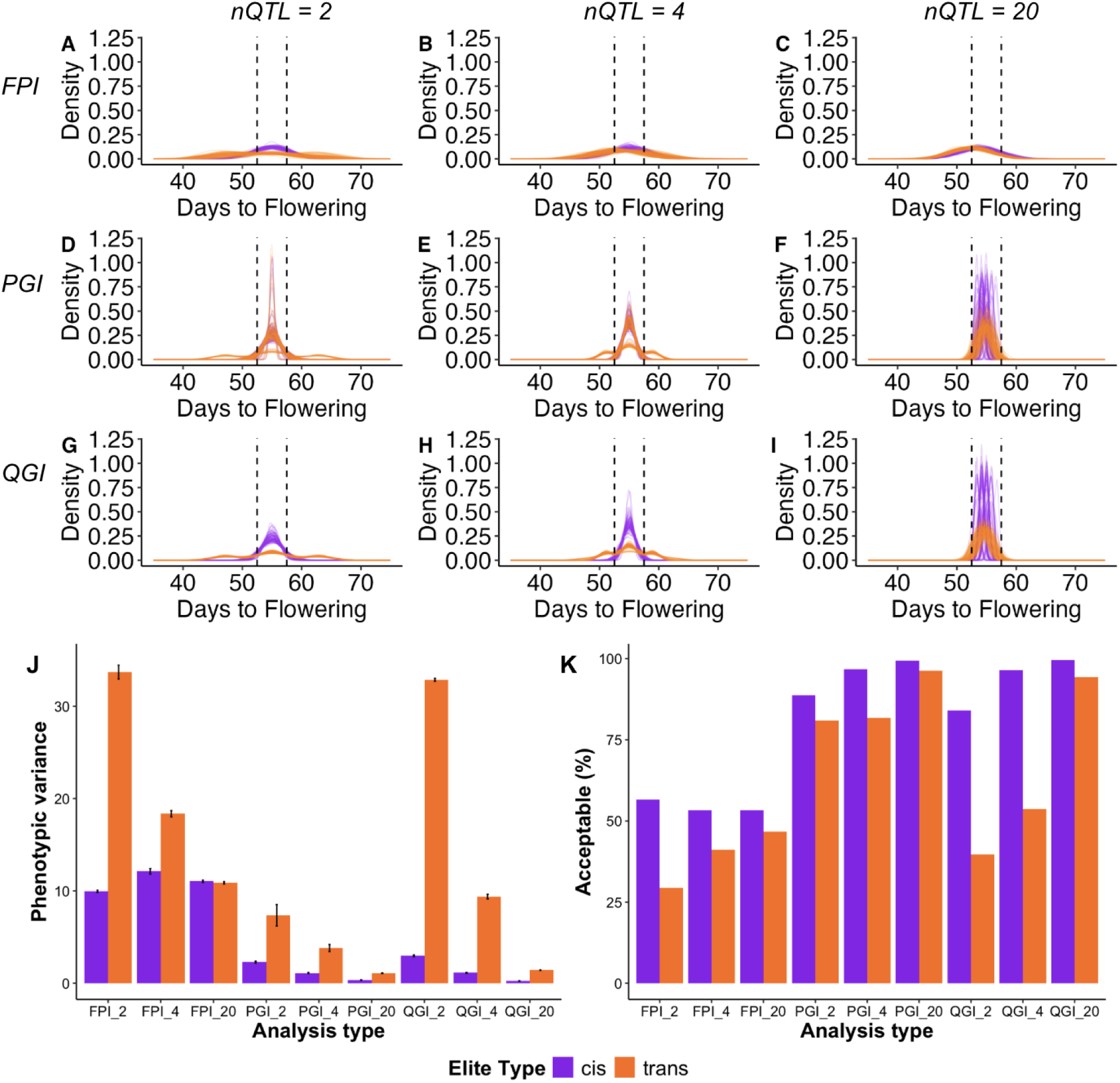
QTL-based genotypic inference (QGI) is the most accurate approach for distinguishing *cis* and *trans*-elite relationships. Phenotypic distributions of F_5_ progenies were compared under three alternative inference approaches: Family-based phenotypic inference (FPI) (panels A–C), QTL-based genotypic inference (QGI) (panels D–F), and Population-based genotypic inference (PGI) (panels G–I). Each panel represents results under different genetic architectures defined by the number of QTLs: 2, 4, or 20. Purple lines represent the distribution of progeny from *cis*-elite crosses (expected to be narrow), while orange lines represent trans-elite crosses (expected to be broader). Average phenotypic variance (J) and acceptable percentage (K) for the simulated acquired trait (flowering time) in F_5_ progeny was computed for each approach (FPI, PGI, QGI) and QTL (2, 4, 20) scenario, labeled as: FPI_2, FPI_4, FPI_20, PGI_2, PGI_4, PGI_20, and QGI_2, QGI_4, QGI_20. For each scenario, 500 individuals were phenotyped, and the phenotypic variance was averaged across 100 simulation replicates.

### Population-based genotypic inference can be effective for polygenic acquired trait architecture

Next we determined the accuracy of using the PGI approach. PGI failed to accurately detect *trans*-elite relationships at a lower number of loci (Figures 5D–E). For QTL numbers of two and four, F_5_ progeny from *trans*-elite crosses showed a narrow distribution for some runs (Figure 5D for QTL = 2; Figure 5E for QTL = 4). Many families of inferred *trans*-elite crosses have narrow distributions leading to erroneous conclusions and thus unpredictable distribution in subsequent generations. In contrast, putative *cis*-elite crosses gave the expected distribution across all 100 runs (Figure 5D for QTL= 2; Figure 5E for QTL = 4). The accuracy of inference for *trans*-elite relationships improved as the number of QTL underlying the acquired trait increased (Figure 5F). At twenty QTL for the acquired trait, *trans*-elite combinations were detected as accurately as *cis*-elite pairs and the accuracy of PGI was similar to QGI (Figure 5F).

### QTL-based genotypic infers elite type across a range of trait genetic architectures

To determine whether QGI in known QTLs can be used to accurately infer *cis-* versus *trans-*elite pairs, we analyzed the phenotypic distribution in F_5_ generations for three genetic architecture scenarios. The progeny of *cis*-and *trans-*elite pairs inferred using QGI follows this prediction (Figures 5G–I). Across all three genetic architecture scenarios, *cis-*elite crosses produce offspring with a narrow distribution for the acquired trait (flowering time in days from sowing to flowering), almost all in the acceptable range of phenotypes. In contrast, *trans*-elite crosses generated offspring with a broad phenotypic distribution for flowering time when this trait is considered to have oligogenic architecture (Figure 5G for QTL = 2, Figure 5H for QTL = 4, and Figure 5I for QTL = 20). In fact, when fewer high-effect variants segregate, the many possible allelic combinations give rise to a wider range of phenotypes for the trait. Thus, most of the progeny from these *trans-*elite crosses are outside the acceptable range of phenotypes and the number of loci is fixed to twenty (Figure 5I). When the acquired trait has multiple small effect variants (nQTL = 20), the phenotypic distribution is still broader compared to the progeny of the *cis-*elite crosses. However, 94% of the offspring remain within the acceptable range of phenotypes as small-effect allele deviates the recombinants less from the initial average (Figure 5K).

### QTL-based genotypic inference provides highest accuracy for elite-type inference

To assess the accuracy of elite-type inference, we compared phenotypic variance within progeny from *cis*- and *trans*-elite crosses across the three approaches (FPI, PGI, and QGI) under simulated genetic architectures involving 2, 4, and 20 loci (Figure 5J). A reliable inference approach is expected to yield low phenotypic variance among *cis*-elite progeny and high variance among *trans*-elite progeny. FPI produced the highest overall phenotypic variance across all scenarios, with limited distinction between *cis*- and *trans*-elite crosses (Figure 5J). While *trans*-elite crosses showed higher variance than *cis*-elite crosses at 2 and 4 loci, the variance among *cis*-elite progeny was unexpectedly high. At 20 loci, variance in *cis*-elite crosses remained elevated, suggesting that FPI becomes increasingly unreliable as trait complexity increases. PGI performed better than FPI, revealing significant differences in variance between *cis*- and *trans*-elite crosses (*p*-values: 0.0015, 0.0131, and 0.0007 for 2, 4, and 20 loci, respectively) (Figure 5J). However, the magnitude of variance separation was modest compared to QGI. QGI consistently outperformed the other approaches, showing highly significant differences in phenotypic variance between *cis*- and *trans*-elite progeny at all locus levels (*p* < 0.001) (Figure 5J). It also demonstrated a clear trend: variance decreased with increasing locus number for both cross types, but remained distinctly lower for *cis*-elite and higher for *trans*-elite crosses.

### QTL-based genotypic inference is the most cost-effective approach when trait loci are known

We compared the cost-effectiveness of two genotypic inference approaches PGI and QGI against the FPI approaches, assuming prior knowledge of trait-associated loci (Table S3). When the number of loci was low (≤4), QGI was the most cost-effective, with estimated costs of $75 and $85 per line for 2 and 4 loci, respectively, compared to $125 for PGI. However, QGI costs rose with increasing locus number and surpassed PGI when 20 loci were involved ($165 per line). In contrast, when trait loci were unknown and required discovery via genome wide association studies (GWAS) or linkage mapping, QGI became significantly more expensive $843 and $670 per line, respectively exceeding both PGI and FPI (Table S4). Under comparable assumptions, FPI, which relies on crossing and phenotyping, incurred an estimated cost of $904 per line per environment (Table S3). Overall, PGI emerged as the most cost-effective approach when no prior knowledge of trait loci was available.

### Genetic gain under recurrent selection is more rapid for *cis*-elite crosses

To evaluate the impact of *cis*-versus *trans*-elite crossing strategies on genetic gain for a desired trait, we simulated 10 cycles of phenotypic recurrent selection, evaluating the performance of F_5_ progenies. We initially hypothesized that *cis*-elite crosses, despite potentially compromising genetic diversity for the acquired trait, would still match or exceed genetic gain in the desired trait (e.g., yield potential). However, the relative performance of elite gene pool crossing strategy depended strongly on the subpopulation origin of the parental lines.

For the acquired trait, progenies from *cis*-elite crosses maintained a stable population mean and narrow phenotypic variance across all cycles, indicating strong trait stabilization (Figure 6A). *Trans*-elite progeny exhibited broader variation early on but gradually stabilized (Figure 6B), while total genetic variance declined over time for both strategies, with *trans*-elite crosses showing an initial peak due to recombination (Figure 6C). For the desired trait, outcomes differed based on whether *cis*-elite parents originated from the same or different subpopulations. When *cis*-elite parents shared the same subpopulation, *trans*-elite crosses yielded greater genetic gain over time (Figure 6D–F). Although *trans*-elite progenies started with broader phenotypic variance (Figure 6E), this variance declined over cycles and contributed to a stronger selection response. Genetic variance for the desired trait was higher in *trans*-elite crosses during the early cycles, supporting this advantage (Figure 6F). However, when *cis*-elite parents originated from different subpopulations, they outperformed *trans*-elite crosses in final genetic gain (Figure 6G–I). These crosses combined subpopulation diversity while retaining elite backgrounds, achieving both trait stability (Figure 6G) and higher final population means (Figure 6H), with comparable variance reduction over time (Figure 6I).

**Figure 6.**
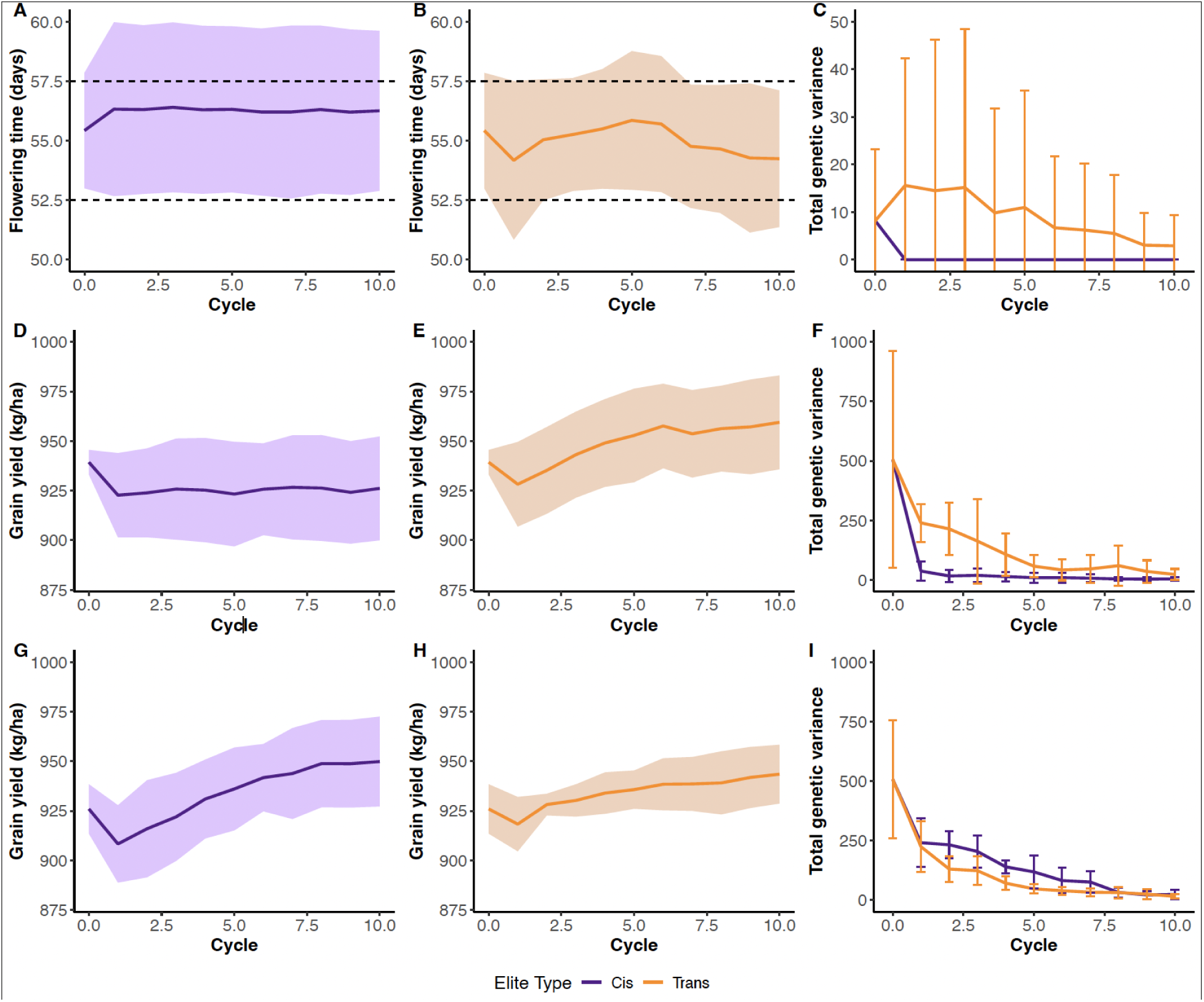
Genetic gain from *cis*-elite crosses surpasses *trans*-elite, but only when parents originate from different subpopulations. Average population mean and total genetic variance were tracked across selection cycles for progeny derived from *cis*- and *trans*-elite crosses. Panels A–C: Results for an oligogenic acquired trait, (A) population mean from *cis*-elite crosses, (B) population mean from *trans*-elite crosses, and (C) total genetic variance over cycles (*cis* vs. *trans*). Panels D–F: Results for a polygenic desired trait and *cis*-elite parents from the same subpopulation, (D) population mean from *cis*-elite crosses, (E) population mean from *trans*-elite crosses, (F) total genetic variance over cycles (*cis* vs. *trans*). Panels G–I: Results for a polygenic desired trait and *cis*-elite parents from the different subpopulations, (G) population mean from *cis*-elite crosses, (H) population mean from *trans*-elite crosses, and (I) total genetic variance over cycles (*cis* vs. *trans*).

## DISCUSSION

### A modernized theoretical and applied framework for eliteness in breeding programs

Many nascent breeding programs, particularly in developing countries, are implementing individual-based phenotypic selection (IPS; Table 1, Figure 3C) in crosses of locally-adapted landrace-derived varieties, but seeing little success in terms of genetic gain or varietal adoption (Walker & Alwang, 2015). Breeding efforts in low-income resource contexts have historically relied on phenotypic selection without access to genomic tools, so lines with locally-adaptive acquired traits are often designated as *elite* based on phenotypic performance alone. Our findings show that crosses between such lines can produce unexpected segregation patterns, indicative of underlying genetic heterogeneity (Figure 1, 2, 4). These outcomes suggest hidden genetic heterogeneity and contradict the notion that elite status equates to genetic homogeneity, a premise often overlooked or taken for granted in breeding literature (Falk, 2010) (Table S1). To address this gap, we develop a framework for defining and operationalizing *eliteness* in nascent breeding populations. By explicitly separating these two categories of traits (acquired versus desired), breeding strategies can be better aligned to harness recombination potential for desired traits while maintaining necessary levels of stability for acquired traits (Figure 2, 6). While we developed the theory and approaches with a focus on nascent programs in developing countries, similar genetic heterogeneity may exist within established networks of public programs, such as for winter wheat in the US Great Plains (Ayalew et al., 2020; Tessele et al., 2025), or even mature commercial hybrid maize programs (Technow et al., 2021).

**Table 1.**
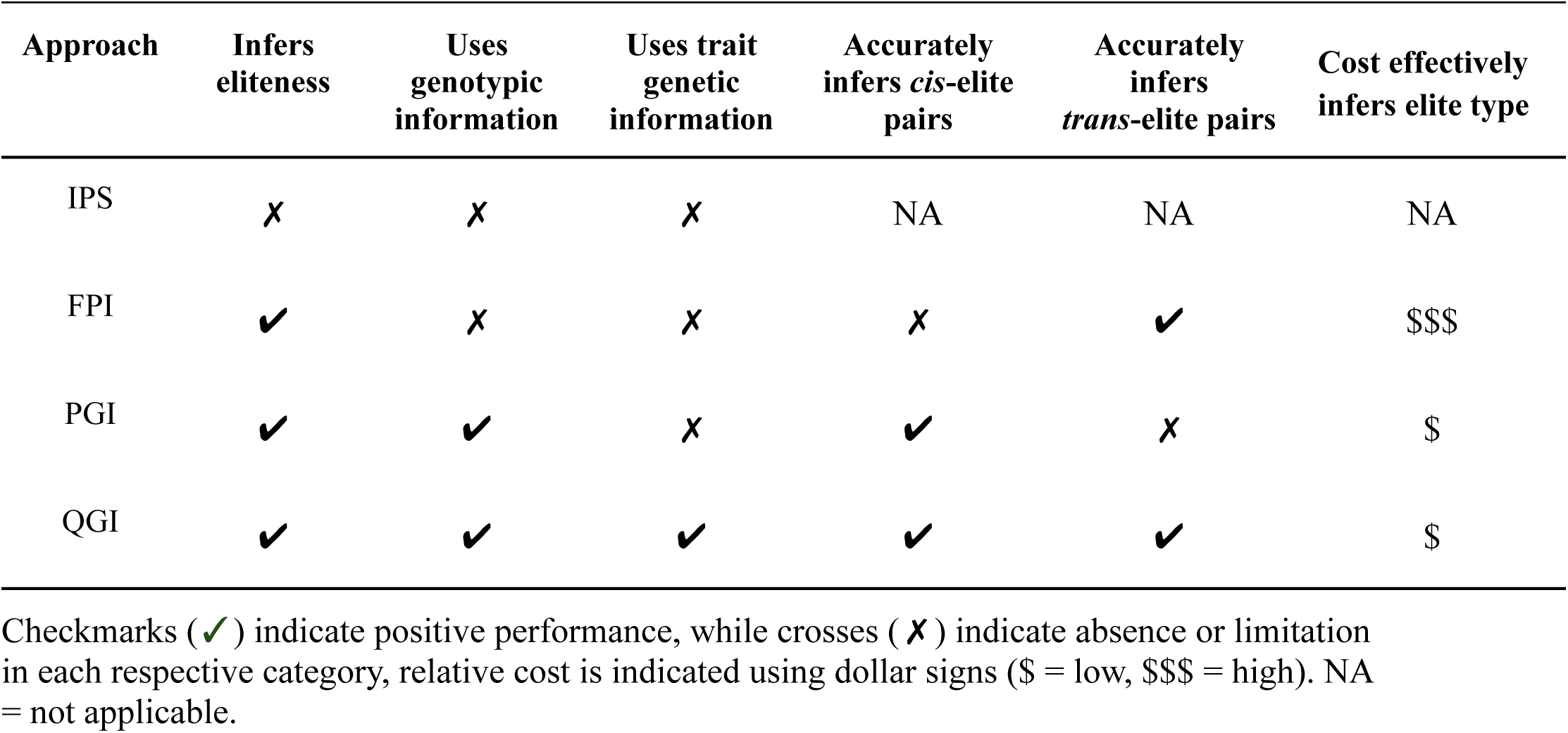
Comparative analysis of breeding approaches based on key evaluation criteria.

Family selection remains a fundamental strategy in plant breeding, particularly in more mature programs where elite lines are extensively tested through their progeny performance (Acquaah, 2015). The theoretical basis of this approach assume that phenotypic differences among families can be attributed to both genetic divergence and the compatibility of parental lines (Walsh & Lynch, 2018). The evaluation of FPI sought to clarify the extent to which these assumptions hold under varying genetic architectures, particularly in distinguishing between *cis*- and *trans*-elite relationships (Figure 5). The rationale behind FPI aligns with classical family selection theory: that phenotypic distributions in progeny reflect underlying genetic structure, with narrower distributions expected from genetically similar (*cis*-elite) crosses, and broader segregation arising from genetically distinct (*trans*-elite) combinations (Falconer, 1996; Walsh & Lynch, 2018b). However, our results reveal a disconnect between this theoretical expectation and empirical outcomes (Figure 5). Across simulated trait architectures ranging from 2 to 20 QTLs, FPI failed to consistently differentiate elite types based on phenotypic variance alone. This finding underscores a key limitation of conventional family-based inference namely, its sensitivity to genetic complexity and its dependence on phenotypic thresholds that may not fully capture the genetic basis of trait expression.

Our findings build on previous theoretical work emphasizing the interplay between within- and between-family genetic variance in shaping selection responses (Fehr, n.d.; Simmonds, 1996; Walsh & Lynch, 2018), and call attention to the need for more robust approaches that integrate genomic information with phenotypic evaluation to identify compatible genotypes. Incorporating a breeder-defined cutoff (i.e., ≥25% of F_2_ individuals outside an acceptable range of phenotypes) improved FPI’s predictive utility to 70% accuracy in oligogenic scenario, suggesting a moderate practical value when trait phenotype is well characterized and thresholds are carefully chosen (Figure 5). However, as the number of QTLs increased, predictive accuracy declined, highlighting the challenges of applying simple inference models to complex, polygenic traits. Interestingly, while we modeled additive acquired traits, the fact that they are under stabilizing selection (i.e. there is a concave trait-fitness function) means that the acquired trait loci are epistatic in terms of fitness (i.e. the fitness effect of an allelic substitution depends on the alleles at other loci). Therefore, one interpretation of our findings is that *cis*-eliteness represents genotypes on the same epistatic peak of an adaptive landscape while *trans*-eliteness represents genotypes on a different epistatic peak (Wade, 1992; Wright, 1932).

### Use genomics findings to guide *de novo* breeding elite gene pools development

The relevance of the QGI approach in this study is closely tied to the nature of the acquired trait, which is controlled by a small number of major-effect loci. This aligns with longstanding debate in crop improvement regarding the value of trait mapping. While some have argued that mapping efforts offer limited utility for improving highly quantitative traits (Bernardo, 2008; Rutkoski, 2019), and that genome-wide selection may yield faster gains (Bernardo & Yu, 2007), our findings suggest that when the architecture of an acquired trait is oligogenic or monogenic, trait mapping is both relevant and impactful. Under such conditions, QGI targeting known loci offered high accuracy, biological relevance, and cost-effectiveness for elite type inference (Figures 5D–F, Table 1). These findings reinforce the value of causal gene identification in enhancing breeding efficiency, particularly in stress-resilience contexts where traits like flowering time or photoperiod sensitivity are key (Guitton et al., 2018).

Programs wishing to deploy QGI could leverage extensive trait mapping in orphan crops such as sorghum, pearl millet, and cowpea, which are targeted by many emerging breeding programs across sub-Saharan Africa and South Asia. In these crops, acquired traits like flowering time, plant height, and drought tolerance have been successfully mapped due to rich phenotypic variation and segregation within traditional landraces (Boukar et al., 2016; Faye et al., 2021; Haussmann et al., 2002). Cloned loci such as *Ma1* (SbPRR37) and *Dw1* in sorghum (Murphy et al., 2011; Yamaguchi et al., 2016), and QTLs identified for flowering time and plant height in pearl millet (Varshney et al., 2017) and cowpea (Lo et al., 2018), could be used in QGI to define *cis*-elite gene pools. The QGI approach could leverage this trait knowledge to robustly distinguish *cis*- and *trans*-elite relationships, offering speed and cost advantages that are essential for adoption by resource-limited breeding programs.

Our simulations of PGI provide qualified support for earlier proposals to use population structure to identify breeding gene pools from locally-adapted landraces and global breeding germplasm (Faye et al., 2021). While this strategy is sound when polygenic trait alleles are already fixed (Figure 2B), our results show that PGI performs poorly for oligogenic traits (Figures 5D–E), where trait loci are few and may not align with principal genomic patterns. PGI’s reliance on overall genomic relationships makes it vulnerable when causal variation is confined to specific regions. Additionally, the accuracy of PGI was influenced by the clustering strategy DAPC performance varied based on the number of PC axes selected, a well-known issue that can lead to false subpopulation detection (Cullingham et al., 2023). Although we used the conservative *optim.a.score* method, this can overestimate dimensions under moderate-to-high gene flow. In scenarios where only a few QTLs underlie the trait, such false structure could obscure true elite-type relationships. Nevertheless, as QTLs represent a small fraction of genome-wide data used for clustering, the limitations of PGI under oligogenic control are consistent with theoretical expectations.

By contrast, QGI demonstrated superior performance by targeting genomic regions linked to specific trait loci. It was robust across trait architectures and retained accuracy even when PGI failed (Figures 5G–I). Though PGI and QGI converged in performance for polygenic traits, QGI’s specificity made it a more reliable inference tool overall. Notably, QGI maintained high inference accuracy for monogenic and oligogenic acquired traits, provided that trait loci had been mapped (Figures 5D–F). However, as the number of loci increases, inference precision can decline (Figure 5J), due to the increased number of plausible haplotypes and variation in IBS scores across genome regions (Henden et al., 2018; Waples et al., 2019). IBS-based inference assumes SNPs are in or near causal loci; thus, QGI’s effectiveness depends on accurate trait mapping and marker resolution (Clark et al., 2013). In summary, our findings suggest that molecular marker tools can effectively guide elite gene pool development in emerging breeding programs, especially when the genetic architecture of key traits is well understood. QGI, in particular, stands out as a cost-effective and biologically grounded strategy for inferring *cis*- and *trans*-elite relationships especially when targeting a small number of high-priority traits. As breeding programs continue to face urgent demands with limited resources, approaches like QGI offer a practical path toward accelerating crop improvement.

Finally, our simulations demonstrate that the genetic diversity and the performance of *cis*-elite crosses depend strongly on the subpopulation origin of the parental lines. When *cis*-elite parents are drawn from the same subpopulation, they exhibit reduced genetic variability, which limits transgressive segregation for genetic yield potential and the likelihood of generating superior phenotypes for desired traits (Figure 6D–F). This finding is consistent with prior findings that crossing closely related lines restricts segregational variance, leading to reduced long-term genetic gain particularly when the trait architecture is oligogenic and polygenic (Bernardo, 2020; Clark et al., 2013). In contrast, *cis*-elite crosses between parents from different subpopulations retain the elite background while introducing meaningful genetic divergence, thereby increasing recombination opportunities and enhancing selection response (Figure 6G–I). This suggests that such *cis*-elite crosses combine stability for acquired traits with the genetic divergence typically targeted for desired traits. *Trans*-elite progenies, while initially more variable (Figure 6E), also benefited from increased genetic variance early in selection, which enabled greater short-term gains in cases where *cis*-elite parents were genetically uniform. These findings emphasize that population structure within elite pools is a critical design consideration for maximizing genetic gain under recurrent selection (Falconer, 1996; Swarup et al., 2021).

### Conclusion

This study highlights that presumed elite lines in emerging breeding programs often harbor hidden genetic heterogeneity, leading to unexpected transgressive segregation when crossed. By distinguishing acquired from desired traits, we demonstrate that elite-by-elite crosses vary in outcome depending on the genetic relationship of parents. While costs will vary across programs and over time, in our initial estimates QTL-based genotypic inference (QGI) emerged as the most accurate and cost-effective method for identifying *cis*- and *trans*-elite relationships, especially when trait loci are known. Our simulations show that *cis*-elite crosses between parents from different subpopulations strike the best balance maintaining stability in acquired traits while promoting recombination for desired traits thus optimizing long-term genetic gain. These findings provide a practical extension to the breeder’s equation, showing how informed elite cross design can enhance selection efficiency without increasing population size.

## MATERIALS AND METHODS

### Simulating diverse locally-adapted landraces

The MaCS software (Chen et al., 2009), implemented via the AlphaSimR package (Faux et al., 2016; Gaynor et al., 2021; Gorjanc et al., 2016), was used to generate an ancestral founding population of 500 outbred lines, setting the species history to GENERIC. This parameter enables the simulation of haplotype sequences shaped by historical changes in population size (Faux et al., 2016; Gaynor et al., 2021). Specifically, the effective population size was modeled to change over time: starting at 10,000 individuals 100,000 generations ago and decreasing stepwise to 6,000, 1,500, 500, and finally 100 individuals in the present, thus mimicking periods of reduced genetic diversity. The genome was modeled with 7 chromosomes, consistent with pearl millet (2n = 2x = 14 chromosomes), and each chromosome had 1,000 random segregation sites. The physical and genetic lengths were set to 1×10⁸ base pairs and 1 Morgan, respectively. Haplotype sequences were generated using the Markovian Coalescent Simulator (MaCS), with a mutation rate of 2.5×10⁻⁸ per base pair (Chen et al., 2009). The founding population was simulated independently 100 times using the same parameters to mimic 100 independent breeding programs (Figures 3 and 4).

For each simulation, two categories of traits were defined in the founding population: a moderately heritable acquired trait (e.g., flowering time or plant height), and a lowly heritable desired trait (e.g., genetic yield potential). These traits were modeled based on phenotypic values observed in the Senegalese pearl millet local germplasm. The acquired trait represented by flowering time was simulated with a mean of 55 days, variance of 10 days, and narrow-sense heritability (h²) of 0.6. The acceptable phenotypic range for flowering time was defined by setting minimum and maximum values at 52.5 and 57.5 days, respectively, to guide stabilizing selection. The desired trait grain yield was modeled with a mean of 900 kg/ha and a low heritability of 0.1, reflecting its complexity and high environmental influence in landraces. Grain yield potential for each individual was modeled based on its phenotypic value for the acquired trait, with penalties applied to those whose trait values fell outside the acceptable range of phenotypes. The further an individual’s phenotype deviates from the optimal mean and acceptable range for flowering time, the greater the penalty applied to its grain yield potential.

Three oligogenic genetic architecture scenarios were simulated for the acquired trait, assigning 2, 4, or 20 quantitative trait loci (QTLs). In contrast, the desired trait was modeled as polygenic, controlled by 100 QTLs. All QTLs were independently and randomly distributed across the chromosomes. For each QTL scenario, equal additive effect sizes were simulated using an external function, defined by the following equations (see Figures 3 and 4).

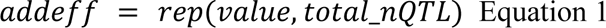

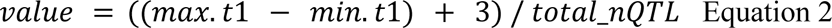

Where total_nQTL is the number of loci controlling the trait, max.t1 and min.t1 represent the maximum and minimum values of the acceptable range of phenotypes. These equations ensure that the additive effects are scaled to maintain trait variability across different QTL numbers.Five subpopulations of 100 individuals each were derived from an ancestral founder population (Figure 3). Within each subpopulation, individuals underwent random mating for 50 generations. At each generation, the top 10% of individuals based on grain yield and falling within an acceptable range for flowering time were selected and recycled for the next round of mating. This selection and mating scheme was applied independently within each subpopulation, mimicking local adaptation to similar environmental conditions while allowing genetic drift to shape allele frequencies over time.

### Simulating purification of landraces to create initial local varieties

Following 50 generations of selection and random mating within each of the five subpopulations, the resulting lines were merged to form a single assembly germplasm representing a broad base of locally adapted diversity. From this assembly germplasm, 200 individuals were selected to represent distinct landrace lines, capturing both phenotypic stability in the acquired trait and variation in the desired trait. These selected lines formed the founding breeding population for subsequent improvement. To generate initial local varieties suitable for selection and further evaluation, each of the 200 lines was advanced through selfing using the single seed descent (SSD) method, producing homozygous elite lines. This purification step served to stabilize trait expression and create a panel of fixed lines representing locally adapted landraces with elite potential.

### Simulating cross-breeding of initial varieties without eliteness inference

To simulate conventional improvement strategies without prior inference of elite relationships (individual-based phenotypic selection; IPS), the top 10% of highest-yielding lines among 200 purified lines were designated as elite. Trait donor lines were selected from within this elite group, each carrying beneficial alleles for specific agronomic traits of interest (e.g., drought tolerance, disease resistance). A forward breeding approach was employed: selected local varieties were crossed with these trait donors, and the resulting progeny underwent phenotypic screening to identify individuals demonstrating superior performance for both introduced and target traits. Progeny exhibiting favorable phenotypes were advanced through standard breeding pipelines, with top-performing individuals selected as candidates for multi-location and advanced field trials. This strategy reflects typical breeding practices in the absence of prior elite-type inference and served as a comparative baseline for evaluating the effectiveness of structured gene pool approaches.

### Phenotypic inference of elite gene pools

FPI was deployed using a half-diallel mating design to evaluate all possible combinations among selected elite parental lines. The F_2_ progenies derived from each cross were assessed to characterize the distribution of phenotypic variation within families. These distributions were used to infer the relationships (*cis* vs. *trans*) and define gene pools. Crosses that produced F₂ populations with narrow phenotypic distributions were inferred to involve parents in a *cis* relationships, while those with broader distributions were interpreted as *trans* relationships, reflecting greater segregation variance. To validate these initial inferences, we examined the distribution of phenotypic variation within F_5_ progenies derived from the same crosses.

### Genotypic inference of elite gene pools

For QGI, we extracted the genotype information in the QTL and then calculated the identity By State (IBS) for each pair of individuals for each marker position using the cluster function daisy() from the R package (Figure S3). Pairs of individuals with the highest IBS indices equal to 1 were considered *cis* elite pairs (Figure S3). In contrast, pairs of individuals with the IBS indices inferior to 1 were considered *trans* elite pairs (Figure S3).

For PGI, a data matrix of molecular markers across all segregating sites was used to first identify clusters using successive K-means with the function find.clusters(). A first discriminant analysis of principal components (DAPC) was performed with these clusters using the dapc() function and then used optim.a.score() to define the optimal number of Principal Components (PCs). A second DAPC was performed using the defined optimal number of PCs. Euclidean distance was used to infer *cis*- and *trans*-elite relationships based on clustering (Figure S3). The Euclidean distance between individuals between and within clusters using the following equation:

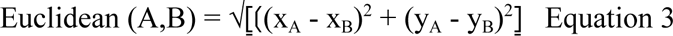

The pairs of individuals with the lowest and highest Euclidean distance within the cluster with the highest proportion of successful reassignment were inferred as having *cis-*elite and *trans-*elite relationships, respectively (Figure S3).

### Analysis of the phenotypic distribution of offspring

Five pairs of putative *cis-* and *trans-* elite combinations were crossed respectively to sequentially generate one F_1_ progeny and selfed to obtain 100 F_2_ progeny then, 100 F_5_ individuals per single-seed descent from each pair (Figures 3 and 4). The phenotypic distribution for flowering time of F_5_ progeny of *cis-* and *trans-* elite crosses was analyzed using a histogram in *ggplot2* R package, respectively. The accuracy of elite type prediction was therefore assessed based on the phenotypic distribution. While a narrow distribution is expected for F_5_ progeny of *cis-* elite crosses, a narrow distribution is expected for F_5_ progeny of *trans* elite crosses (Figures 1C-D).

### Comparing costs of de *novo* gene pool strategies

We estimated the costs of defining breeding gene pools for 50 elite lines using QGI and PGI and compared them to FPI. Cost estimates were based on the millet breeding program budget in Senegal during the 2016/2017 season (Table S2). Figure S1 outlines the sequence of steps considered for each strategy. The phenotypic approach involved crossing the 50 elite lines with four required control lines (depending on the trait) using a half-diallel mating design. F_1_ seeds were harvested from each cross and then advanced to F_5_ generation progenies, following the procedures previously described (Figure 4). The phenotypic distribution of the F_5_ progenies was then analyzed for the trait of interest.

For the genotypic approaches, we considered two scenarios: (i) major QTLs already known, and (ii) QTLs unknown and requiring discovery. The latter scenario includes additional steps such as forward genetic methods GWAS and biparental QTL mapping as well as reverse genetic approaches like allele mining and comparative genomics supported by functional genomics (Figure S1). In this study, cost estimates were based solely on forward genetics. For GWAS, we estimated the cost of genotyping and phenotyping a diversity panel of 300 lines across two sites over two years. For biparental QTL mapping, we simulated the development and evaluation of 250 F_2_ individuals, including genome-wide genotyping and trait phenotyping. Once major QTLs were identified, the next steps included genotyping the 50 elite lines at these specific regions for allele screening and computing similarity matrices for QGI analysis. To estimate PGI costs, we assumed genome-wide genotyping of the 50 elite lines followed by PGI analysis for elite clustering.

Genotyping costs were estimated at $20 per line for genome-wide data and $5 per line for each QTL-specific assay based on previous genotyping activities conducted in Senegalese the sorghum breeding program. Phenotyping costs were estimated at $64 per line, based on the same 2016/2017 Senegalese pearl millet program budget. The experimental design used to derive phenotypic cost estimates was an alpha-lattice design (7 × 13), with three replications. Each elementary plot consisted of two rows of 8 mounds, flanked by two border rows, with 90 cm spacing between rows and between mounds. Costs included all field operations in a rainfed environment from land preparation to harvest covering labor, inputs, and equipment (Table S2).

### Simulating intercrossing to initiate recurrent selection program

To initiate the recurrent selection program, three additional acquired traits were simulated, resulting in a total of four acquired traits and one desired trait modeled in the ancestral founding population. The purification process to create initial local varieties was conducted as previously described. QGI was used exclusively to infer *cis-* and *trans-*elite relationships among parental lines. These inferred elite parent pairs were then used to initiate a phenotypic recurrent selection (PRS) program. In cycle 0 (C0), two scenarios were simulated independently: one starting with five *cis-*elite pairs and the other with five *trans-*elite pairs. From each cross, one F_1_ plant was generated and selfed to produce 100 F_2_ individuals via single seed descent, resulting in a total of 500 F_5_ individuals per scenario (Figure 4). These F_5_ individuals were advanced through subsequent generations via the same method. At each cycle, selection was based on performance for the desired trait (e.g., grain yield), while maintaining acquired traits within an acceptable range of phenotypes. This applied directional selection on the desired trait and stabilizing selection on the acquired traits. The phenotypic traits were modeled as independent traits in the simulation. A total of ten selection cycles were simulated for each scenario to evaluate the long-term effectiveness of the phenotypic recurrent selection strategy. The outcomes of the nascent breeding program were evaluated by tracking phenotypic trends for both acquired and desired traits across selection cycles using *ggplot2* in R. In addition, total genetic variance for these traits was extracted and visualized to assess the impact of selection on genetic diversity over time.

## ACKNOWLEDGEMENTS

This study is made possible through funding by the Feed the Future Innovation Lab for Collaborative Research on Sorghum and Millet through grants from American People provided to the United States Agency for International Development (USAID) under cooperative agreement number AID-OAA-A-13-00047 and 7200AA-19LE-00005. The contents are the sole responsibility of the authors and do not necessarily reflect the views of USAID or the US Government. Support was also provided by the Gates Foundation through the grant “Green Evolution - Accelerating Dryland Cereals Improvement for Africa (INV-053669)” to GPM.

---

**Table S1.**
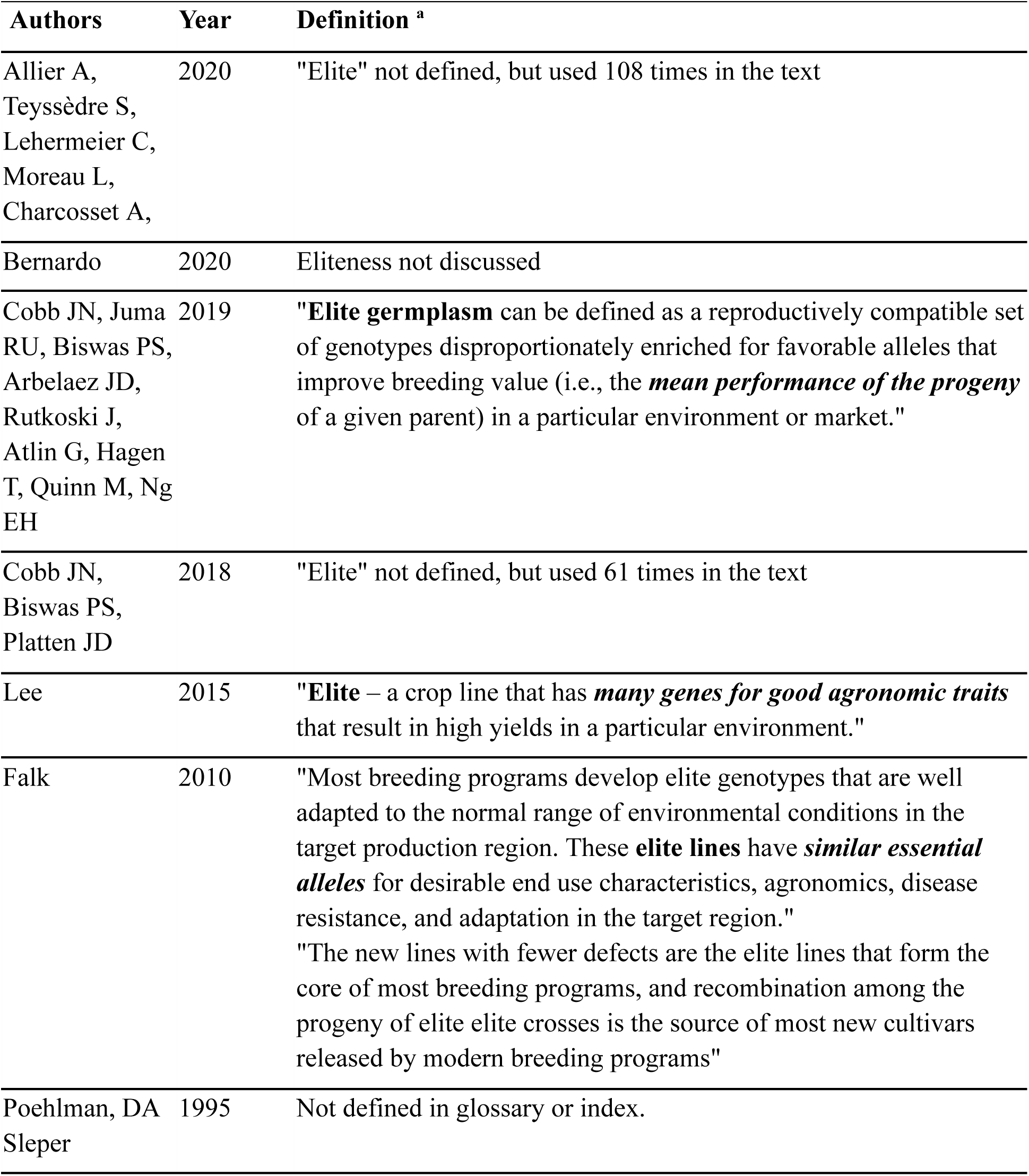
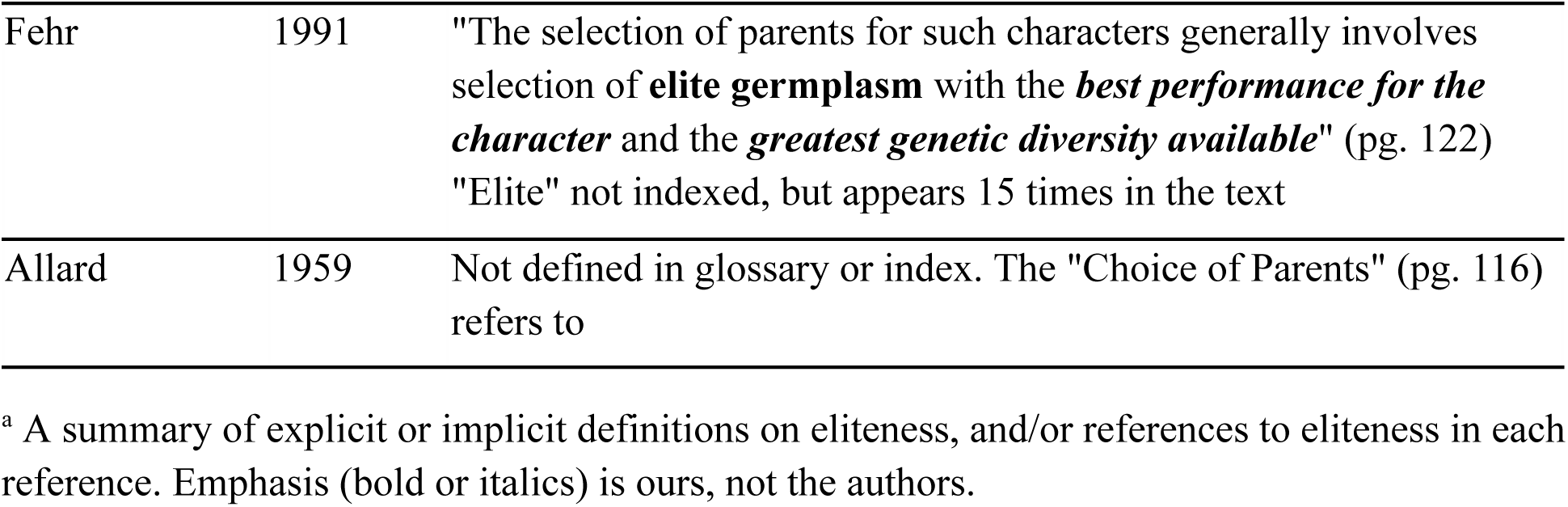
Conceptual ambiguity in the definition of *eliteness* in representative contemporary and classical plant breeding literature.

**Table S2.** Estimated field phenotyping budget per cycle for 404 pearl millet lines from the national core collection at the ISRA research station in Bambey, Senegal.

|  | **Activities included** | **Cost (FCFA)** | **Cost (USD)** |
| --- | --- | --- | --- |
| 1. Field preparation | Plowing | 25,000 | \$42 |
| 2. Inputs | Fertilizers, pesticides, and others products | 150,000 | \$250 |
| 3. Labor | Temporary labor, Daily labor | 3,200,000 | \$5,333 |
| 4. Material and supplies | Small material, Bags, Labels | 200,000 | \$332 |
| Total | - | 3,525,000 | \$5,875 |
| **Total per line** | - | **38,315** | **\$63** |
| **Total per plant** | - | **8,725** | **\$22** |

Costs are presented in both the original currency (CFA Franc) and in US dollars, assuming a conversion rate of 1 USD = 600 FCFA.

**Table S3.** Estimated per line costs of phenotypic and genotypic screening strategies for characterizing de novo elite gene pools in a nascent breeding program.

| **Analysis <sup>a</sup>** | **Cost (USD/line)** |
| --- | --- |
| FPI | \$904 |
| QGI, assuming 4 known QTL | \$85 |
| QGI with linkage mapping to identify QTL | \$670 |
| QGI with GWAS to identify QTL | \$843 |
| PGI | \$125 |

<sup>a</sup> FPI (Family-based Phenotypic Inference): Relies on replicated phenotyping across environments; cost is per line per location.

**Table S4.** Estimated screening costs for QTL-based genotypic inference (QGI) across varying numbers of known QTL.

| **Analysis <sup>a</sup>** | **Cost (USD) <sup>b</sup>** |
| --- | --- |
| QGI_2 | \$75 |
| QGI_4 | \$85 |
| QGI_20 | \$165 |

<sup>a</sup> Numbers indicate QTL count per scenario.
<sup>b</sup> Costs assume prior knowledge of QTL and are based on the use of KASP markers for genotyping.

